# Challenges and Efficacy of Astrocyte-to-Neuron Reprogramming in Spinal Cord Injury: In Vitro Insights and In Vivo Outcomes

**DOI:** 10.1101/2024.03.25.586619

**Authors:** Alessia Niceforo, Lyandysha V. Zholudeva, Silvia Fernandes, Yashvi Shah, Michael A. Lane, Liang Qiang

**Author notes:** Corresponding authors; Correspondence to. Alessia Niceforo, PhD / Liang Qiang, PhD Department of Neurobiology & Anatomy Drexel University, 2900 W Queen Lane Philadelphia, PA, 19129; /. Co-first author.

## Abstract

Traumatic spinal cord injury (SCI) leads to the disruption of neural pathways, causing loss of neural cells, with subsequent reactive gliosis and tissue scarring that limit endogenous repair. One potential therapeutic strategy to address this is to target reactive scar-forming astrocytes with direct cellular reprogramming to convert them into neurons, by overexpression of neurogenic transcription factors. Here we used lentiviral constructs to overexpress *Ascl1* or a combination of microRNAs (miRs) *miR124, miR9/9\**and *NeuroD1* transfected into cultured and *in vivo* astrocytes*. In vitro* experiments revealed cortically-derived astrocytes display a higher efficiency (70%) of reprogramming to neurons than spinal cord-derived astrocytes. In a rat cervical SCI model, the same strategy induced only limited reprogramming of astrocytes. Delivery of reprogramming factors did not significantly affect patterns of breathing under baseline and hypoxic conditions, but significant differences in average diaphragm amplitude were seen in the reprogrammed groups during eupneic breathing, hypoxic, and hypercapnic challenges. These results show that while cellular reprogramming can be readily achieved in carefully controlled *in vitro* conditions, achieving a similar degree of successful reprogramming *in vivo* is challenging and may require additional steps.

## Introduction

The injured adult mammalian spinal cord is incapable of significant repair. This limitation is due in part to two major neuropathological consequences of spinal cord injury (SCI): i) the destruction of neuronal connectivity, loss of spinal neurons and cystic cavitation (Lane, White et al. 2008) and ii) the subsequent formation of growth-inhibitory fibroglial ‘scar’ surrounding the lesion site, which includes activated astrocytes (Lang, Cregg et al. 2015). To advance regenerative medicine and effectively repair the damaged spinal cord, it is crucial to develop reliable methods for efficiently generating new populations of neural cells. Cellular reprogramming holds the potential to address this challenge. This rapidly evolving technology involves the use of transcription factors to directly convert one cell type into another (El Wazan, Urrutia-Cabrera et al. 2019). Takahashi and Yamanaka first developed this technology by successfully reprogramming somatic cells into induced pluripotent stem cells (Takahashi and Yamanaka 2006). This pioneering work established cellular reprogramming as a transformative approach for generating patient-specific cells (Hung, Khan et al. 2017), and paved the way to new strategies that can directly reprogram cells (Bajohr and Faiz 2020). Direct reprogramming allows rapid conversion between two somatic cell types without laborious and prolonged cloning procedures, and without the de-differentiation process associated with increased risk of tumorigenesis. This technique mitigates the need for more invasive delivery of cells via cell transplantation and subsequent immunosuppression (Zhang, Chen et al. 2022) (Xu, Du et al. 2015). The wound healing/scarring and proliferation of glial cells in response to neural injury represents an endogenous source of cells that might be harnessed with reprogramming methods to create new populations of neurons after SCI.

A plethora of studies have indeed validated direct conversion from somatic cells into functional neurons, utilizing various sets of master neurogenic transcription factors. More recently, some of these factors have been tested for therapeutic efficacy in repairing damage to the central nervous system (CNS) (Vignoles, Lentini et al. 2019) (Pereira, Birtele et al. 2019, Pereira, Birtele et al. 2019) (Clark, Roman et al. 2022). Advances have been achieved through delivery of reprogramming constructs into the brain, which have shown some benefit, but still require further validation (Parmar and Pereira 2021, Vasan, Park et al. 2021, Bocchi, Masserdotti et al. 2022) Indeed, the optimal transcription factor(s) to drive conversion, and means of promoting their expression, remain poorly defined. Furthermore, it has been challenging to differentiate between glial cells successfully reprogrammed into neurons and resident neurons that were inadvertently infected by viral vector injection. Finally, the current strategies to assess functional recovery still pose challenges in separating the contributions of reprogramming and repair from those of spared tissue and plasticity.

Here we employed two different lentivirus-based reprogramming approaches, each targeting a distinct signaling pathway involved in CNS development. We tested these approaches *in vitro* in both cortically-derived and spinal cord-derived astrocytes. Consistent with previous reports, cortically-derived astrocytes were effectively converted to cells that expressed neuronal proteins and exhibited electrophysiological patterns consisted with mature neurons. In contrast, conversion of spinal cord-derived astrocytes was less effective with these reprogramming factors. Additional testing was performed *in vivo* in adult rats after cervical SCI to assess whether the approach could be more effective in the lesion environment where activated astrocytes accumulate. Although limited, this *in vivo* reprogramming promoted a degree respiratory functional recovery of suggesting the therapeutic potential of this approach. Contrary some prior published studies, however, and consistent with the results from *in vitro* studies, *in vivo* reprogramming of astrocytes in the injured spinal cord was less successful than initially predicted.

The result from this work suggest that no single reprogramming method can be used to reprogram cells, even the same cell type (astrocytes) from similar tissues (the central nervous system).

## Results

### Direct in vitro reprogramming efficiencies for astrocytes isolated from neonatal brain

Given that *Ascl1* (Addgene, #27150), and the combination of *miR124, miR9/9\** and *NeuroD1* (miR124,miR9/9*+NeuroD1, Addgene, #31874, named as “miRs+ NeuroD1” from here on), have each demonstrated efficiency in converting different types of somatic cells into functional neurons, we initially tested their capacity to convert activated astrocytes into functional neurons *in vitro*. We first used the Tet-on lentiviral system to introduce either Ascl1 cDNA or miRs+ NeuroD1 into primary astrocyte cultures derived from either neonatal (postnatal day 2-3) cerebral cortex, before subjecting them to an *in vitro* differentiation protocol (Figure 1A).

**Figure 1.**
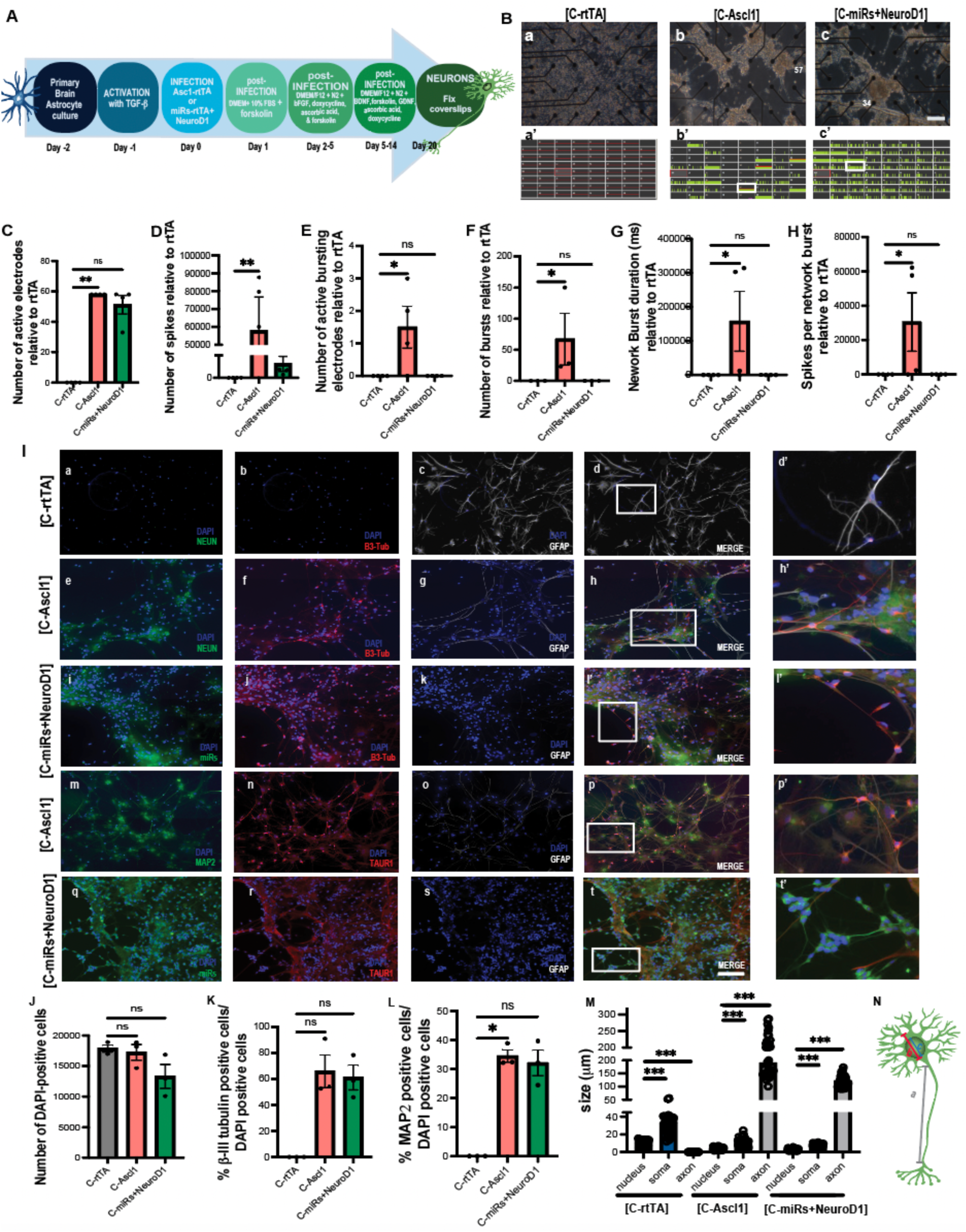
I*n vitro* conversion of cortical astrocytes. (A) Timeline of *in vitro* experimentation. Primary astrocyte cultures were generated from postnatal day 2-3 (P 2-3) rat cortex, that were activated with TGF-ß 1 day (day -1) prior to viral transduction (LentiGFAP:Ascl1, LentiGFAP:miR124+miR9/9*+NeuroD1, day 0). Infection was removed day 1 after viral transduction. Then, on day 2, doxycycline was added to the media, allowing neuronal conversion Cells were fixed and stained at day 20 of reprogramming. (B) Experiments conducted to measure field potential activity using the multi-electrode array (MEA). (Ba, Bb, Bc) Phase images of re-plated cells on MEAs on Day 20 of reprogramming. Scale bar: 250 microns in Ba,Bb,bc. (Ba’, Bb’, Bc’). Pictures of activity recorded around each electrode with green lines representing spikes. Note absence of spikes in control (C-rtTA) compared to reprogrammed neurons (C-Ascl11 and C-miRs+NeuroD1). Green marks =neuronal spikes; red bars =neural bursts. The number of corresponding electrodes and the respective recording cells were indicated in the phase contrast images and highlighted in b, b’ and c, c’. (C) Graph shows the average number of active electrodes per well. (D) Bars show the number of spikes in active electrodes per well. (E) Graph shows the average number of actively bursting electrodes per well with at least one burst recorded. (F) Graphs show the number of bursts relative to the control (rtTA). (G) The graph shows the network burst duration (ms). The number of spikes per burst were showed in H. (I) Immunocytochemistry for astrocytic marker GFAP, neuronal nuclear protein NeuN and neuronal cytoskeletal marker β-III tubulin were used to assess protein expression *in vitro*. (Ic) The bright white spots indicated that C-rtTA is positive for GFAP, while the lack of green and red stain meant negative for NeuN and β-III tubulin. (Id’) Magnification and display detail of image Id. (Ig) The faint white stain for GFAP in C-Ascl1-treated cells indicated some positivity for astrocytes, however the increase in NeuN (De) and β-III tubulin (If) labeling supports conversion of these treated cells. (Ih’) Magnification and display detail of image Ih. (Ii) Green signal indicates the infected cells with miRs+NeuroD1. (Ik) Low GFAP reactivity for miRs+NeuroD1 was indicative of low number of astrocytes. (Ij) The positivity for β-III tubulin along the processes was indicative of a high neuron population. (Il’) Magnification and display detail of image Dl. Scale bar: 250 microns in Ia-It. (J) Cell viability (percent of DAPI-positive cells) at day 20 of reprogramming. (K-L) Bar graphs show quantification of the percent of B3 tub-positive cells per total DAPI-stained cells (K), and the percent of MAP2-positive cells per total DAPI-stained cells (L) at day 20 of reprogramming. (M) Relationship between neurite length, nuclei and soma size for each condition C-rtTA, C-Ascl1and C-miRs+NeuroD1 at day 20 of reprogramming. (N) Schematic representation of the parameters considered in M (a=axon length, b=soma, c=nucleus). All data are represented as mean ± SEM normalized to vehicle per line (C-rtTA). Analyses were conducted using one-way ANOVA. *p<0.05; **p<0.005; ***p<0.0001; ns=not significant.

Cortical astrocytes treated with *Ascl1* or *miRs+ NeuroD1* (C-Ascl1 and C-miRs+ NeuroD1, respectively) were compared with control-treated cells that received *rtTA* (reverse tetracycline-controlled transactivator, C-rtTA,). At the beginning of reprogramming (day -2), the cortical astrocytes showed a typical star-shaped morphology (Supplementary Fig. 1A, a). This morphology changed after the TGF-β treatment (day -1, Supplementary Fig. 1A, b) and after infection (day 0, Supplementary Fig. 1A, c). From day 1 to day 5, elongation of cellular processes was observed across all groups (Supplementary Fig. 1A, e,h,k), with the most pronounced effect seen in the Ascl1 and miRs+ NeuroD1-treated cells. Particularly, at day 20 of reprogramming, cells that received C-Ascl1 and C-miRs+ NeuroD1 exhibited a typical neuronal morphology, characterized by elongated cell body, extension of neurites and dendrites, and soma formation (Supplementary Fig. 1A, i and l), and showed minimal astrocytic features whereas brain astrocytes treated with rtTA retained characteristic astrocytic morphology (Supplementary Fig. 1A, f).

Immunohistochemistry supported these findings (Fig. 1I). At day 20 of reprogramming, control-treated cells (C-rtTA) were positively labelled with antibodies to GFAP (glial fibrillary acidic protein; Fig. 1D, c), while negative for the neuronal nuclear protein (NeuN; a.k.a. Fox3) (Fig. 1I, a), and β-III tubulin (Fig. 1D, b). In contrast, cortical cells treated with Ascl1 and miRs+ NeuroD1 were positive for both β-III tubulin and NeuN (Fig. 1I, e,f, j), and negative for GFAP (Figure 1I, g, k). Consistent with their neuronal identity, C-Ascl1 and C-miRs+ NeuroD1 treated cells were also positively labeled for microtubule-associated protein 2 (MAP2) and the total tau marker (TAUR1) (Fig. 1I, m,n,r).

To assess cell death and survival following lentiviral infection, the number of cells surviving after treatment was quantified relative to the number of cells plated at the start of the experiment (20,000 cells), for each specific treatment. At day 20, the final day of differentiation, approximately 83% of Ascl1-treated cells persisted. In contrast, only 65% of cells infected with miRs+ NeuroD1 survived (Fig. 1J). The proportion of neuronal (β-III tubulin-positive) cells was also measured relative to the number of DAPI-positive cells (Fig. 1K). Although no β-III tubulin-positive cells were observed in C-rtTA treated cells, 65% and 58% of cells were β-III tubulin-positive in the C-Ascl1 and C-miRs+ NeuroD1 groups, respectively. Thus, both survival and reprogramming efficiency was marginally better in the C-Ascl1 treated cells. A consistent reduction in the ratio of β-III tubulin to total DAPI positive cells was found in C-miRs+ NeuroD1 (Fig. 1K).

Next, we used MAP2 to quantify the proportion of mature neurons (Fig. 1L). While there were no MAP2-positive cells in the C-rtTA treated cells, 35% and 30% of cells were MAP-2 positive in the C-Ascl1 and C-miRs+ NeuroD1 groups, respectively. Recognizing that the neuronal length assessment is a crucial aspect of determining neuronal maturity (Niceforo, Marioli et al. 2021) (Compagnucci, Piermarini et al. 2016), and understanding the significance of the interplay between neuron length, nucleus size, and soma size in each neuronal cell (Tramontin, Smith et al. 1998, Thompson and Brenowitz 2005) we quantified these parameters across all conditions (Fig. 1M). These analyses revealed that cell lines treated with *Ascl1* or *miRs+ NeuroD1* had neurites measuring three times the length of their soma diameter (Fig. 1N). The average length of neurites in treated cells were 188.8µm (±SD= 46.6) and 121.3µm (±SD=21.0) in the C-Ascl and C-miRs+ NeuroD1 groups, respectively.

The functional characteristics of reprogrammed cells were electrophysiologically assessed using multi-electrode arrays (MEAs; Multi Channel Systems 60MEA200/30iR-Ti-gr). The cells were plated at a density of 50,000 cells per well, and spontaneous activity in each group was recorded for 5 minutes on day 20 of the reprogramming (Fig. 1B). Phase contrast images (Fig.1B a,b,c) were used to assess presence and morphology of plated cells, and with C-Asl1 and C-miRs+ NeuroD1 groups showed elongated, neuron-like processes. Cells from these groups also showed spontaneous neuronal spikes and bursts during recording, albeit with a significative difference in the number of spikes between treatment groups (Fig. 1B b’,c’). Control-treated (C-rtTA) cells showed no spontaneous detectable activity (Fig. 1B a’).

While spontaneous activity in cells from C-Ascl1 and C-miRs+ NeuroD1-treated groups showed a substantial number of active electrodes and detectable spikes (Fig.1C-D), only C-Ascl1 treated cells exhibited spontaneous burst activity (Fig. 1E-H). Neuronal activity was evaluated and confirmed pharmacologically by adding synaptic antagonists into the cell culture media. The introduction of AP5 (a selective NMDA receptor competitive antagonist) to the culture medium reduced activity in C-Ascl1 and C-miRs+ NeuroD1 after 20 min, and activity was abolished after 60 minutes (Supplementary Fig. 1C). Conversely, the inclusion of bicuculine (GABAergic antagonist) resulted in increased number of spikes and bursts (supplementary Fig. 1D).

### Reprogramming efficiency in spinal cord-derived astrocytes

Next, we tested the potential of lentiviral transduction of *Ascl1* and *miRs+ NeuroD1* to reprogram spinal cord astrocytes. While the isolation and preparation of the spinal cord astrocytes required slightly different conditions that that of cortical astrocytes, the reprogramming strategy used was comparable (generating SC-rtTA, SC-Ascl1 and SC-miRs+ NeuroD1 groups, respectively; Fig. 2A). Although there was morphological change over time it was distinct from that seen in cortical astrocytes (Supplementary Fig. 2B). Particularly, by day 15 of reprogramming, both SC-rtTA, SC-Ascl1 and SC-miRs+ NeuroD1 cells displayed an atypical morphology characterized by poorly defined extensions (Supplementary Fig. 2B f,i,l). Consistent with these data, immunohistochemistry revealed only partial expression of the neuronal markers β-III tubulin, NeuN, MAP2, and TAUR1 in the SC-Ascl1 and SC-miRs+ NeuroD1 groups (Fig. 2F). In contrast, GFAP reactivity was seen throughout cells in both treatment groups. As expected, the control SC-rtTA group was positively labelled for GFAP, and negative for both β-III tubulin and NeuN (Fig. 2F a,b,c). Notably, survival of spinal cord reprogrammed astrocytes was reduced compared cortically-derived neurons. About 50% of cells treated with *rtTA* remained viable, while the survival of SC-Ascl1 and SC-miRs+ NeuroD1 cells was approximately 20% (Fig. 2G). Furthermore, only 10% of those surviving cells were positively labelled for β-III tubulin and MAP2 (Fig. 2H-I). While SC-rtTA cells showed no evidence for neurite extension, short neurites were measured in SC-Ascl1 and SC-miRs+ NeuroD1 groups (Fig. 2J) revealing neurite length to be less than 3-times the soma diameter.

**Figure 2.**
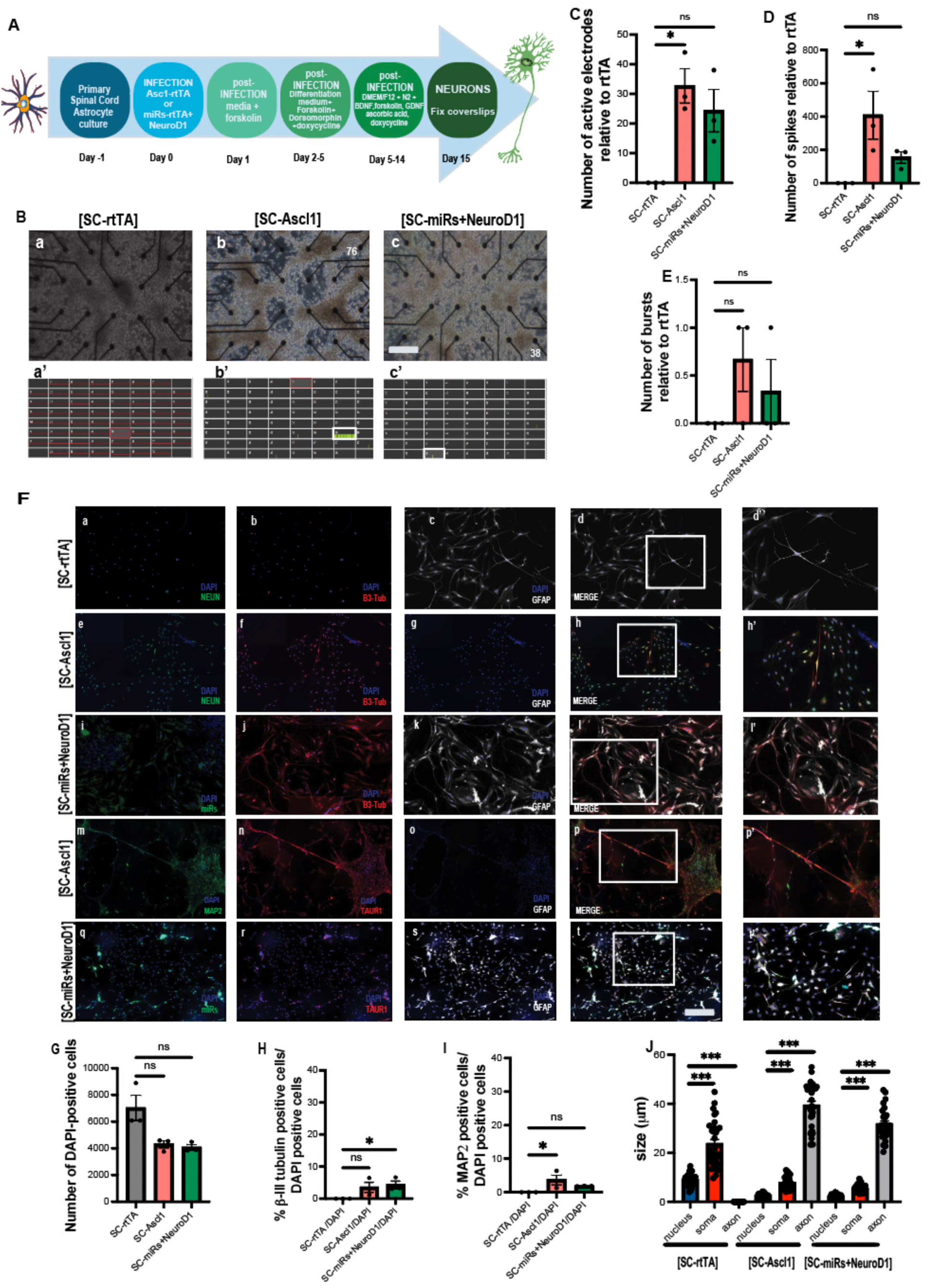
I*n vitro* conversion of spinal cord astrocytes. (A) Timeline of *in vitro* experiment. Spinal cord astrocytes were generated from postnatal day 2-3 (P 2-3), and then infected with lentivirus LentiGFAP:Ascl1, LentiGFAP:miR124+miR9/9*+NeuroD1 at day 0. On day 1 the infection was removed; the post infection media was changed at day 2 and doxycycline was added. (B) Measurement of the potential activity using MEA. (Ba, Bb, Bc). Phase images of re-plated cells on MEAs on Day 15. (Ba’, Bb’, Bc’) Neuronal activity was recorded, and for each condition a total (SC-rtTA) or partial (SC-Ascl1 and SC-miRs+NeuroD1) absence of spikes was observed. Scale bar: 250 microns in Ba,Bb,bc. (C) Graph shows the average number of active electrodes per well. (D) Bars show the number of spikes in active electrodes per well. (E) Graph shows the average number of actively bursting electrodes per well with at least one burst recorded. (F) Spinal cord reprogrammed cells stained for the astrocytic marker GFAP, and neuronal markers NeuN, β-III tubulin, MAP2 and TAUR1. The white signal in Dc indicated that SC-rtTA was positive for GFAP, while the lack of green and red stain meant negative for NeuN (Da) and β-III tubulin (Fb). (Fd’) Magnification and display detail of image Dd. SC-Ascl1 and SC-miRs+NeuroD1 resulted also positive for GFAP (Fg,Fk,Fo,Fs) and only partially positive for β-III tubulin, NeuN, MAP2 and TAUR1 (Fe, Ff, Fj, Fm, Fn, Fr). (Fh, Fl, Fp, Ft) Magnification and display detail of image Fh’, Fl’, Fp,’ Ft’. Scale bar: 250 microns Fa-t. (G) Number of surviving and DAPI-positive cells at the end of reprogramming (day 15). (H-I) Percentage of spinal cord reprogrammed cells positive cells for β-III tubulin and MAP2 relative to DAPI staining at the conclusion of the reprogramming process (day 15). (J) Relationship between neurite length, nuclei, and soma size (μm). All data are represented as mean ± SEM normalized to vehicles per line (SC-rtTA). Analyses were conducted using one-way ANOVA. *p<0.05; **p<0.005; ***p<0.0001; ns=not significant.

Cells from both SC-Ascl1 and SC-miRs+ NeuroD1 groups that were plated on MEAs had short processes (Fig. 2B b,c), and there were no measurable processes in SC-rtTA cells (Fig. 2B a). While there were no detectable spikes in SC-rtTA cells (Fig. 2B a’), some spikes were recorded in SC-Ascl1 and SC-miRs+ NeuroD1 cells (Fig. 2B b’,c’), but fewer than were seen in reprogrammed cortical astrocytes. Therefore, we can say that that the lentivirus-induced reprogramming of spinal cord astrocytes is less efficient than cortical astrocytes.

### *In vivo* astrocyte to neuron conversion was limited

Despite limited evidence for spinal astrocyte-to-neuron reprogramming *in vitro*, the primary goal of these studies was to assess whether reprogramming could be achieved *in vivo* after traumatic SCI. Initially, we injected viral constructs into the intact cervical spinal cord of rats (n=15) to assess risk of inflammation and tissue damage associated with the injection and viral infection. Regardless of the virus used (GFAP;mCherry, GFAP:Ascl1-T2A-mCherry, or GFAP:GFP-miR124+miR9/9*), the intraspinal injections resulted in a small degree of pathology limited to the site of injection (Supplementary Fig. 1A-C). Accordingly, there is greater rationale for viral delivery directly into the lesion cavity of injured animals, which minimizes tissue pathology, while targeting the glia comprising the fibro-glial border of the lesion.

To test the potential for *in vivo* astrocyte reprogramming in the injured spinal cord, we used a rat model of cervical contusion injury, which results in cystic cavitation surrounded by an accumulation of reactive glial cells (contributing to the ‘fibroglial scar’), identified by immunolabeling with antibodies to GFAP (Supplementary Fig. 1D, E). The injury was generated between cervical level 3 and 4 (C3/4) with an Infinite Horizon Pneumatic Impactor (Precision Systems, Lexington, Kentucky) with a force preset to zero dwell time and 200 kD (see Materials and Methods). The anatomical and functional deficits resulting from this injury have previously been characterized (Spruance, Zholudeva et al. 2018) (Zholudeva, Iyer et al. 2018). Two of the 42 animals used for this study did not survive the injury, and one animal died later.

One week after injury, animals (n=39) were anesthetized and the spinal cord re-exposed for sham injection (exposure without injection; n=7), or intraspinal injection of either control (GFAP;mCherry; n=9), Ascl1 (GFAP:Ascl1-T2A-mCherry; n=9) or miRs+NeuroD1 (GFAP:GFP-miR124+miR9/9*; n=8) viral construct into the lesion epicenter (9.2 ±0.8 microlitres). The animals were left to recover for 3 weeks for histological assessment and 8 weeks, at which point they underwent terminal bilateral diaphragm electrophysiology and histological assessment (Fig. 3 A, B).

**Figure 3.**
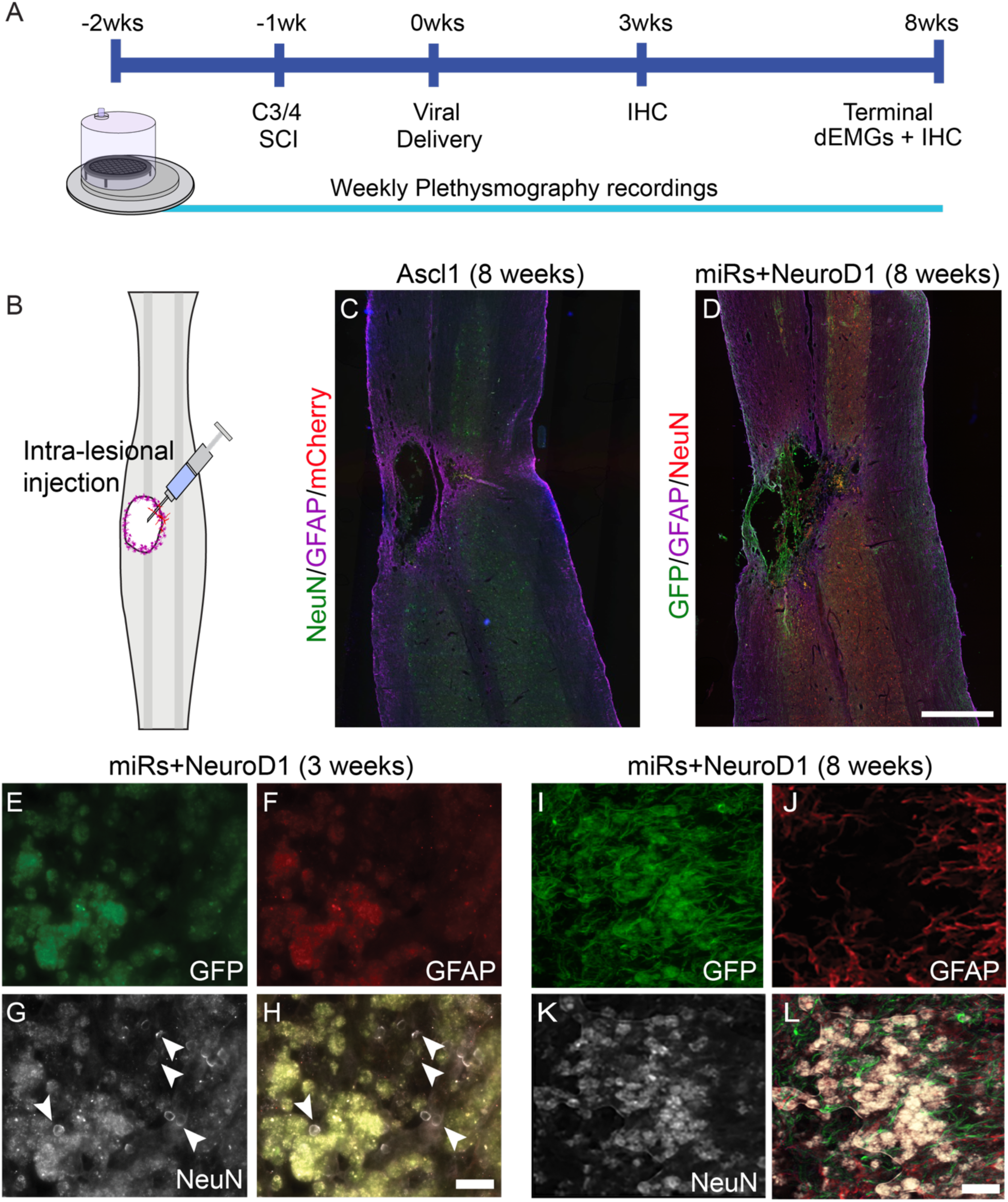
Intraspinal astrocyte-to-neuron reprogramming. Experimental timeline of the *in vivo* strategy for intraspinal astrocyte-to-neuron reprogramming, with plethysmography prior to- and weekly post-contusion injury (A). Viral constructs were injected into the lesion cavity 1-week post-injury (B) and reprogramming was assessed via immunohistochemistry 3 and 8 weeks post-viral delivery. Histological sections (horizontal, 40um) of the cervical spinal cord at the site of injury and viral injections (Ascl1, C; miRs+NeuroD1, D), showing the lesion cavity 8 weeks post-injury, surrounding by infected cells (expressing mCherry (C) or green fluorescent protein (GFP; D)). High magnification images of the lesion cavity sub-acutely after injury (3 weeks after miRs+NeuroD1 injection) immunohistochemically labelled for GFP (E), GFAP (F), NeuN (G) with merged image shown in (H) reveals cells with macrophage morphology. These larger cells are interspersed with smaller NeuN-positive cells with a morphology atypical of mature neurons (G). High magnification images of lesion cavity 8 weeks post-injury and injection (miRs+NeuroD1, I-L) immunohistochemically labelled for GFP (I), GFAP (J), NeuN (K) with merged image shown in (L) reveals absence of smaller NeuN-positive cells, and the cavity comprises putative macrophages only. Scale bar: 500um in C-D and 50um in E-L.

The distribution of the injected virus was determined by either GFP or mCherry fluorescence (from viruses expressed miRs or Ascl1, respectively) following injection directly into the lesion cavity a week post-injury resulted in infection of cells within and immediately surrounding the injury (Figure 3A). In addition to the expression of viral fluorophores (Fig. 3D), sections were immunolabeled with neuronal (NeuN and β-III tubulin) proteins (Fig. 3 E, F), and antibody to astrocytic GFAP (Fig. 3G). Three weeks after injection, cells in both Ascl1 or miRs+NeuroD1-treated animals were positively labeled for fluorescent reporters and expressed neuronal markers but were GFAP-negative (Fig. 3G). These putative neurons had an atypical morphology (Fig. 3 D’, E’, F’, G’), however, and were limited primarily to the lesion epicenter and were densely surrounded by macrophages. While their appearance was not comparable to maturing neurons seen throughout the spinal cord, they closely resembled reprogrammed neurons in the brain that have been described in previous studies (Gotz and Bocchi 2021, Bocchi, Masserdotti et al. 2022) (Heinrich, Bergami et al. 2014). By 8 weeks post-injection, there was no evidence for these putative reprogrammed cells in any tissues examined and labeled proteins were restricted to the vesicles of infiltrating macrophages within the lesion cavity (Fig 3 H,I,J,K arrows). GFAP immunolabeling was examined in serial sections (30µm thick, 90µm apart) in each animal. GFAP-positive glial reactivity around the lesion cavity was comparable between all animals (Supplementary Figure 1).

### Phrenic and respiratory function after contusive cervical SCI and *in vivo* reprogramming

Next, we used whole-body plethysmography to assess respiratory behavior in control, Ascl1- and miRs+NeuroD1-treated animals respectively, in baseline (normoxic), hypercapnic and recovery conditions. Patterns of breathing under baseline and hypoxic conditions were not significantly affected by the treatments. Both breathing frequency and tidal volume during hypercapnic challenge were reduced acutely (during the first week) after injury in all groups but recovered to pre-injury level by two weeks after injury (data not shown). These changes, which were comparable across all control and treated animals, recovered to pre-injury levels by two weeks after injury. There was no significant difference between the control, Ascl1 and miRs+NeuroD1 treated animals. This pattern of functional plasticity and recovery is consistent with prior assessments of ventilation after mid-cervical contusion injury (Lane, Lee et al. 2012).

Terminal, bilateral diaphragm electromyography (dEMG) was used to assess functional respiratory outcomes in injured animals, 8 weeks after viral injection. Integrated diaphragm activity (Fig. 4A, top trace) derived from raw output (Fig. 4A, bottom trace) was used to compare average diaphragm amplitude (Fig. 4B, see Methods for details) during eupneic breathing (Fig. 4B, black trace), hypoxic challenge (Fig. 4B, blue trace) and hypercapnic challenge (not shown). A significant difference in average diaphragm amplitude was seen between Ascl1 and miRs+NeuroD1 recipients during eupneic breathing (Fig. 4C), hypoxic (Fig. 4D), and hypercapnic (Fig. 4E) challenges, with miRs+NeuroD1 recipients having higher average amplitude than Ascl1-recipients. The ability of each animal to respond to a respiratory challenge (i.e., the difference between baseline diaphragm amplitude and amplitude seen during a respiratory challenge, Fig. 4 A, B, blue line) was also quantified. While no differences were observed in responses to hypoxic challenge (Fig. 4F), miRs+NeuroD1 recipients on average showed a greater response than Ascl1-treated animals (Fig. 4G).

**Figure 4.**
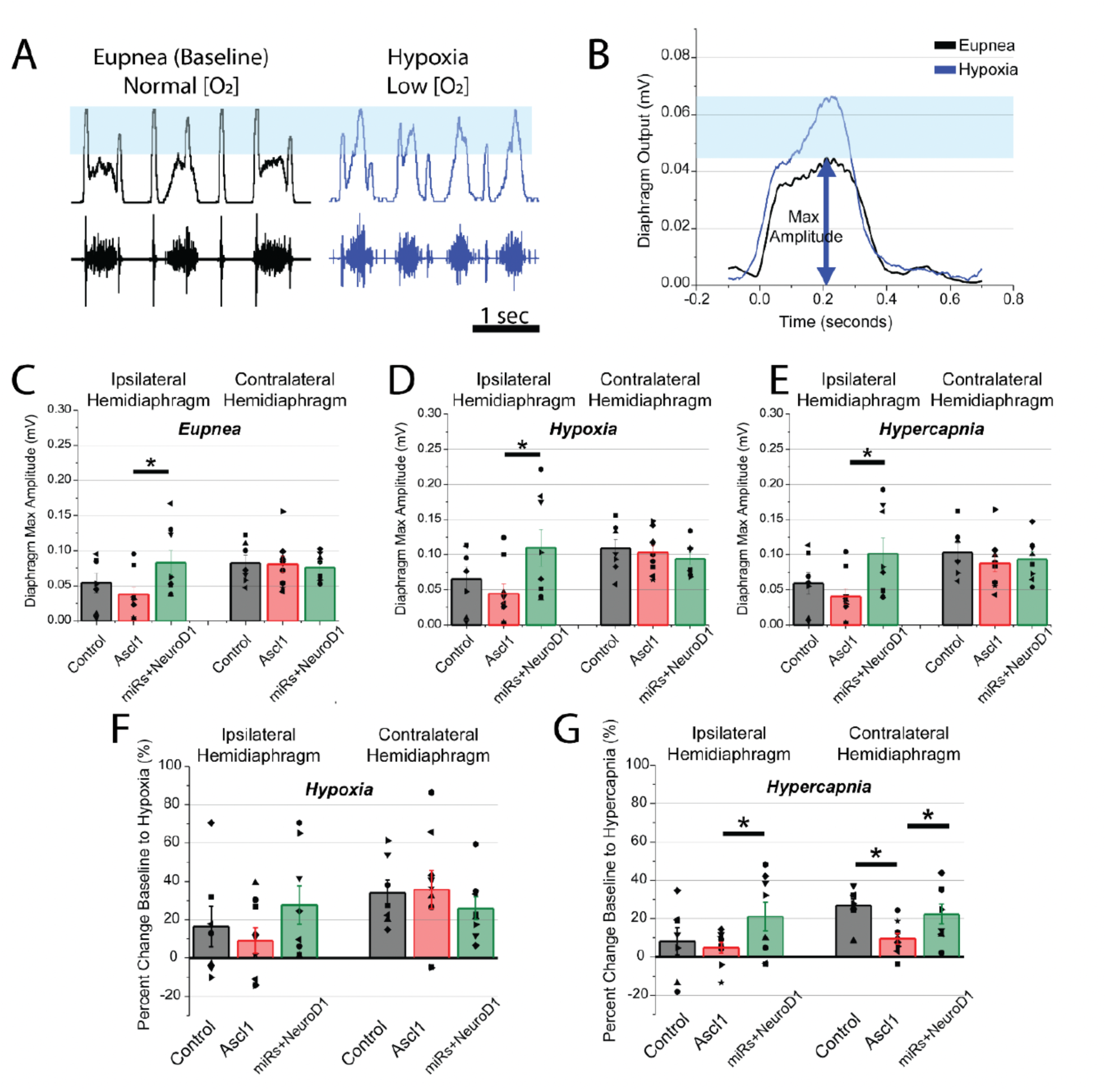
Bilateral electromyography of diaphragm activity reveals modulation of output. (A) Representative traces of muscle output of ipsilateral-to-injury hemidiaphragm (bottom trace) and integrated signal (top trace) during eupneic breathing (black traces, left) and hypoxic challenge (blue traces, right). (B) Average traces of integrated activity spanning 40 seconds of the recording during eupneic breathing (black trace) and hypoxic challenge (blue trace) were used to quantify the average maximum amplitude and response to challenge (difference between maximum diaphragm amplitude during eupnea and respiratory challenge, blue box). (C-E) Quantitative analysis of average diaphragm output during eupneic (C) or challenged breathing (hypoxia in D and hypercapnia in E) reveals significant difference between Ascl-1 and miRs+NeuroD1 recipients. (F-G) Percent change baseline to hypoxia (F) and hypercapnia (G) reveals a difference between Ascl-1 and miRs+NeuroD1 recipients during hypercapnic challenge. Each dot represents an individual animal’s average diaphragm output over a 40 second time span. All data are represented as mean ± SEM; *p≤0.05, student t-test.

## Discussion

Cervical SCI is characterized by significant neuronal loss and phrenic network degeneration (Baussart, Stamegna et al. 2006, Brown, DiMarco et al. 2006). Restoration of breathing capacity is an important goal since respiratory failure is a leading cause of morbidity and mortality in individuals with SCI (Winslow and Rozovsky 2003, Allen, Nichols et al. 2021). Recent studies in regenerative medicine have strived to address these challenges by targeting the precise replacement of lost neurons and the restoration of the disrupted neural network. While cell transplantation remains a promising approach to address this neural repair (Fischer, Dulin et al. 2020) (Hall, Fortino et al. 2022), reprogramming methods offer a means of harnessing the existing host tissues to produce new populations of cells that are most needed for repair. These approaches include the direct reprogramming of non-neuronal cells into neurons (Gotz and Bocchi 2021). A pivotal scientific advancement in this field was the demonstration that differentiated non-neuronal cells within the central nervous system (e.g., astrocytes), could be directly converted into functional neurons *in vitro* (Heins, Malatesta et al. 2002). Since then, *in vitro* direct reprogramming of astrocytes into neurons has provided valuable insights into the molecular mechanisms underlying the conversion and the challenges that hinder efficient reprogramming (Hersbach, Simon et al. 2022). However, comparing *in vitro* and *in vivo* experiments is challenging due to methodological differences of the *in vitro* vs. *in vivo* reprogramming and the distinct environments in which the cells operate (i.e. isolated system *in vitro* vs. complexity of biological systems *in vivo* ). Here, we tested two different strategies: 1) The possible differences of *in vitro* reprogramming of two different types of astrocytes (derived from cortex and spinal cord of postnatal rats) into functional neurons using the transcription factor *Ascl1* and the combination of microRNAs+NeuroD1; and 2) the possibility of conducting *in vivo* reprogramming of astrocytes following spinal cord injury, using a rat contusion SCI model.

Consistent with previous studies, our *in vitro* data confirmed that reprogramming of cortical astrocytes into functional neurons is achievable with a notably high efficiency using *Ascl1* (Rivetti di Val Cervo, Romanov et al. 2017) and microRNAs+NeuroD1 (Zhou, Gu et al. 2015). This was observed at day 20 of reprogramming, by the typical neuronal morphology shown in C-Ascl1 and C-miRs+NeuroD1 cells, as well as expression of neuronal markers (β-III tubulin, NeuN, MAP2 and TAUR1). To further validate their neuronal identity, spontaneous neuronal activity of C-Ascl1 and C-miRs+NeuroD1 cells was recorded and confirmed pharmacologically by adding synaptic antagonists (AP5 and bicuculine) into the cell culture media. It is important to note that the reprogramming of cortical astrocytes to neurons appeared to be more efficient using the transcription factor *Ascl1* than miRs+NeuroD1, in all aspects analyzed. This may be due to the different mechanisms that the transcription factor *Ascl1* and combination of microRNAs (miR124, miR9/9*) and NeuroD1 use to convert of somatic cells into neurons (i.e., the activation of specific pathways and/or negative regulators of gene expression) (Vasan, Park et al. 2021).

In contrast to these results, *in vitro* reprogramming of spinal cord derived astrocytes showed a lower conversion efficiency. Although there was morphological change over time, at day 20 of reprogramming using either Ascl1 or miRs+NeuroD1, cells displayed poorly defined extensions. The survival of reprogrammed spinal cord derived cells was also less than with cortically-derived cells. Only 10% of the surviving cells were positively labeled for the neuronal markers β-III tubulin and MAP2. Notably, some spontaneous neuronal activity was recorded in SC-Ascl1, and SC-miRs+NeuroD1 cells, but it was far less than in reprogrammed cortical astrocytes. These data raise further questions, potentially linked to distinct astrocyte types (cortical and spinal cord), and the potential insufficiency of the reprogramming factors used. Indeed, several studies showed that for the *in vitro* direct reprogramming not only is it important to consider which transcription factors to overexpress for lineage conversion, but also the starting somatic cell source, including its regional identity (El Wazan, Urrutia-Cabrera et al. 2019, Vasan, Park et al. 2021).

Regarding the *in vivo* direct reprogramming, our data imply the feasibility this technique within the injured spinal cord, showing the transformation of astrocytes into putative neurons positively labeled for neuronal markers (NeuN and β-III tubulin), but with an atypical morphology. Despite their appearance which was not typical of maturing neurons seen throughout the spinal cord, the cells were negative for the glial marker GFAP. Notably, 8 weeks post-injection, we observed an extensive infiltration of macrophages surrounding the lesion cavity, and some smaller cells with a morphology not typical of mature neurons. These results are consistent with previous studies (Heinrich, Bergami et al. 2014, Bocchi and Gotz 2020), where the reprogrammed neurons in the brain showed morphological characteristics deviating from those typically observed in mature neurons. Although *in vivo* reprogramming occurred in the spinal cord injury site, the difference in the efficacy of *in vitro* versus *in vivo* reprogramming data was evident. When considering the operation of transcription factors (i.e. *Ascl1*) and miRNAs, particularly *in vivo*, it’s essential to recognize that the determination of cell fate hinges on the balance between inducers and repressors (Yamashita, Shang et al. 2019). Thus, it is important to investigate the role of several negative regulators of neurogenesis (e.g., the p53-p21 pathway and oxidative stress) that can suppress the reprogramming processes (Jiang, Xu et al. 2015, Masserdotti, Gillotin et al. 2015, Gascon, Murenu et al. 2016, Vasan, Park et al. 2021). Addressing the challenges posed by the *in vivo* environment of the damaged spinal cord will be crucial for effective implementation of direct reprogramming as a therapeutic strategy for neural repair in clinical settings.

Over the last 10 years, numerous attempts at *in vivo* reprogramming in SCI have been conducted using the transcription factors NeuroD1 (Puls, Ding et al. 2020) and Ascl1 (Heinrich, Blum et al. 2010, Liu, Miao et al. 2015) with some success. Importantly, previous studies also showed that the injury-induced reactive astrocytes can resemble neural progenitors in terms of their gene expression profiles (Sirko, Behrendt et al. 2013, Gotz, Sirko et al. 2015). This is in line with our *in vivo* results. So, unique molecular interactions may occur during neuronal conversion and produce distinct phenotypes (Chen and Li 2022).

While *in vivo* neuronal reprogramming has been demonstrated as a possible approach for SCI, reports with functional recovery data at the behavioral level are rare (Tai, Wu et al. 2021), and no integration or electrophysiological parameters were reported. Here, we investigated the functional potential of *in vivo* reprogramming by measuring physiological, behavioral, and anatomical parameters. While notable behavioral and electrophysiological responses were lacking in both of our conversion approaches, we made the novel discovery of acute conversion of local astrocytes at the site of the injury into potential neurons, however, they did not persist long-term.

Despite recent advancements made in direct reprogramming in the last few years, numerous challenges have arisen and require attention and resolution. One of the drawbacks of direct reprogramming techniques is the low efficiency and unstable functionality they entail. Indeed, cells derived from direct reprogramming still exhibit significant differences from the authentic target cells in crucial aspects. Notably, efficiency and function frequently correlate with one another. Based on that, some key areas that can enhance both aspects of direct reprogramming have been identified: 1) modifying the factors involved in direct reprogramming; 2) regulating the epigenetic factors and pathways; and 3) improving cellular maintenance (Zhang, Chen et al. 2022, Gerber and Perrin 2024). Collectively, *in vivo* reprogramming is a relatively new field, and many of the underlying mechanisms have yet to be discovered and verified. Our ongoing research highlights a novel potential therapeutic approach for promoting anatomical repair while identifying important limitations to this technology. Future goals will focus on the development of more efficient and persistent conversion systems, opening new avenues for disease modeling and potential regenerative therapies.

## Acknowledgments

AN was supported by the Craig H. Neilsen Foundation (430069). LVZ was supported by the Lisa Dean Moseley Foundation, NIH NINDS F32 NS119348 and CIRM DISC2-14180. LQ was supported by Craig H. Neilsen Foundation (381793). MAL was supported by R01 NS104291, Wings for Life and the Lisa Dean Moseley Foundation.

We thank Dr. Nicholas Kanaan of Michigan State University for providing TAUR1 antibody; Dr. Tara Fortino from University of British Columbia for her contributions and help to the project; Skandha Ramakrishnan from Drexel University for producing the lentivirus for the study; and Dr. Itzhak Fischer for helpful comments in the preparation of the manuscript.

## Author contributions

Dr. Niceforo carried out and analyzed data from the *in vitro* experiments and wrote the manuscript. Dr. Fernandes carried out and analyzed data from the *in vivo* experiments. Yashvi Shah helped with technical support.

Drs. Qiang, Lane, Zholudeva, and Niceforo designed, supervised, and assisted with experiments, data interpretation, and analysis, and revised the manuscript.

All authors edited, read, and approved the final version of the submitted manuscript.

## Supplementary figures

**Supplementary figure 1:**
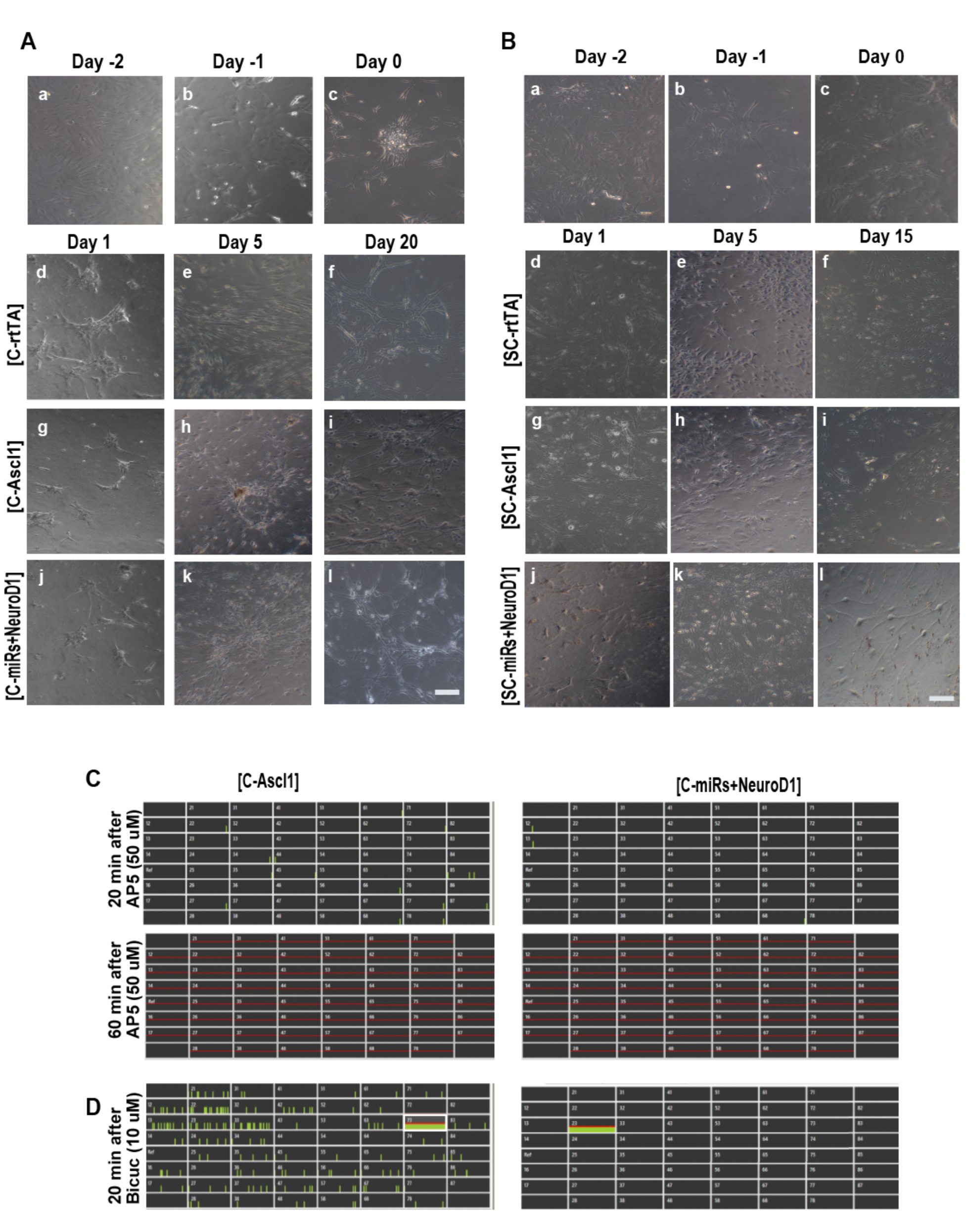
(A) Phase images to investigate changes in cell morphology during conversion of cortical astrocytes. Days -2 through 0 displayed an undifferentiated astrocytic morphology. At Day 1, C-rtTA, C-Ascl1, C-miRs+NeuroD1, promoted elongation of processes, which were greatest in those treated with Ascl1 and miRs. At Day 15, C-rtTA, C-Ascl1, C-miRs+NeuroD1 show further elongation and a large increase in the number of cells/ nuclei. Ascl1 and miRs have clearer neural processes and little visible astrocytic morphology, while rtTA has more astrocytic characteristics. Scale bar: 250 microns in a-l’. (B) Phase images to investigate changes in cell morphology during conversion of spinal cord astrocytes. Scale bar: 250 microns in a-l’. (C-D) Verification of neuronal activity of C-Ascl1 and C-miRs+NeuroD1 through the introducing synaptic antagonists AP5 and bicuculine to the cell culture medium at day 20 of reprogramming. (C) Recordings were conducted 20-60 min after addition of AP5 (50 μM), and 20 min after addition of bicuculline, (10 μM). In C-Ascl1 and C-miRs+NeuroD1, the presence of AP5 in the culture medium led to the decrease (after 20 min) and the complete cessation (after 60 min) of neuronal activity. (D) Conversely, the addition of bicuculline resulted in a heightened occurrence of spikes and burst in both conditions.

**Supplementary Figure 2:**
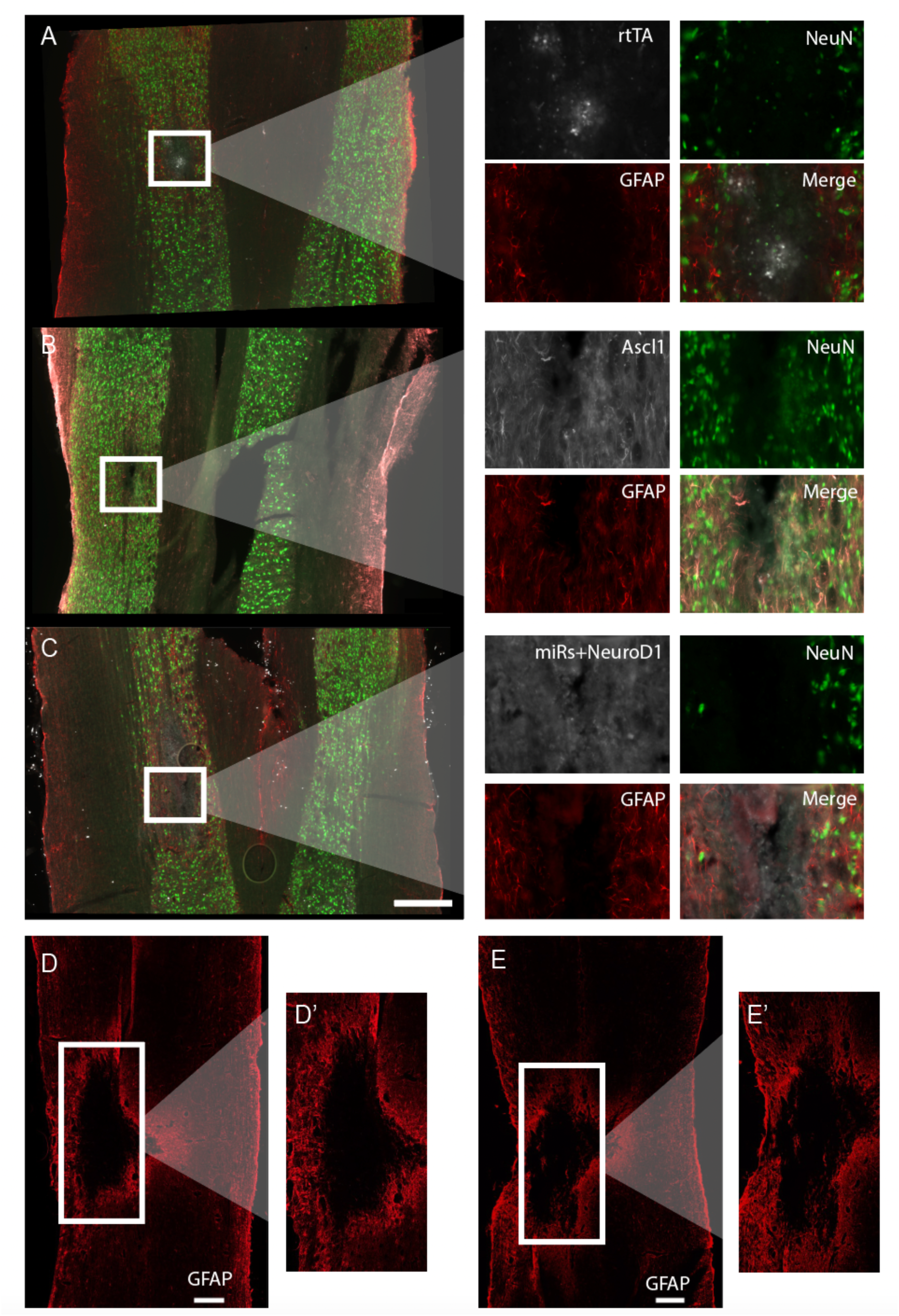
(A,B,C) Longitudinal sections of the naïve spinal cords infected with rtTA (A), *Ascl1* (B), and *miRs+NeuroD1* (C). The spinal cord sections were immunostained with the glial marker GFAP (red), and the neuronal marker NeuN (green). In white the lentivirus used for the infection (rtTA, Ascl1-mCherry and miRs+ NeuroD1-GFP, respectively). The intraspinal injections resulted in small degree of pathology limited to the site of injection. (D, E) Representation of the cystic cavitations in rtTA (D) and Ascl1 (E) after C3/4 contusion. The scars in both the conditions were surrounded by an accumulation of reactive glial cells immunolabeled with GFAP (red). (D’, E’) Zoomed-in image from D and E, respectively. (A, B, C, D, E) Scale bar: 500μm.

## Materials and methods

### Primary Astrocyte culture

Brain primary astrocyte cultures were derived from postnatal day 2-3 (P2-3) neonatal rat cortex. The meninges were removed, the cortex was carefully dissected out, and tissue was dissociated into a single cell suspension by using Trypsin/EDTA 0.25% (Thermo Fischer Scientific, 25200056) and DNase 300 mg/ml (Sigma-Aldrich, 10104159001) and mechanical disruption. The cells were pelleted at 3000rpm for 5 minutes, resuspended, and plated with Minimum Essential Media (MEM) (Thermo Fischer Scientific, 11095072) containing 10% Horse Serum (Thermo Fischer Scientific, 26050088), 0.6% D-(+)-Glucose (Sigma-Aldrich, G8769-100ML), 1% Penicillin /Streptomycin (Thermo Fischer Scientific, 15140122), in T75 flasks coated with Gelatin 0.1% (Sigma-Aldrich, G1393-100ML). Primary cultures of astrocytes were maintained in the incubator at 37°C and 5% CO2 and the culture media was replaced 24 hours later and then every other day. Neuronal and other cell type contamination was avoided by vigorous shaking of the flask. Cells were passaged at 80%–90% confluency using TrypLE™ (Gibco, 12604021). After 3 passages, the cells were frozen in freezing medium consisting of 60% Fetal Bovine Serum (FBS) (Life Technologies, 26140-079), 30% MEM and 10% DMSO (Sigma-Aldrich, C6295-50ML) or fixed to evaluate the purity of the cultures.

To infect the cells *in vitro*, astrocytes were thawed and plated at 20,000 cells/well in MEM with 10% Horse Serum and 6.7% glucose onto 24 well plates (Nest Scientific, 70201) containing 25mm coverslips pre-coated with 1mg/ml of Poly-L-ornithine (Sigma-Aldrich, P4957-50ML) for 24 hours. Astrocytes were activated with 20ng/ml of TGF-ß (R&D Systems, 100B001) suspended in Dulbecco’s Modified Eagle Medium (DMEM) (Corning, 10013CV) with 10% FBS. 24 hours after activation, astrocytes were infected with 1.3µl of either Ascl1 (10^7^ - 10^8^pfu) or miR24+miR9/9*+NeuroD1 (10^7^ - 10^8^pfu) in DMEM with 10% FBS. The media was changed 16 hours after infection to fresh DMEM with 10% FBS and 5μM Forskolin (Sigma-Aldrich, F6886).

From day 2 to day 5 post-infection, the media was replaced with DMEM/F12 (1:1) (Thermo Fisher Scientific, 11330057) with N2 supplement (Thermo Fischer Scientific, 17502001), 10ng/ml basic fibroblast growth factor (bFGF) (Peprotech, 100-18B), 200 μM Ascorbic acid (Sigma-Aldrich, A5960-25G), 5μM Forskolin and 2ug/ml doxycycline (Sigma-Aldrich, D9891-10G). Thereafter, from day 6 to day 15 post-infection, the medium was changed every other day consisting of DMEM/F12 (1:1) with N2 supplement, 0.2M Ascorbic acid, 5μM Forskolin, 2μg/ml doxycycline, 20 ng/ml Brain-derived neurotrophic factor (BDNF) (PreproTech, 450-02), 20ng/ml Glial cell line-derived neurotrophic factor (GDNF) (PreproTech, 450-10) and 5μg/ml laminin (Invitrogen, 23017015) added weekly into the medium.

To obtain spinal cord astrocytes, postnatal (P2-P3) rats were euthanized, and their spinal cord rapidly isolated from the vertebral column. The meninges were removed using the dissecting microscopes (Zeiss Stemi DV4), tissue was centrifuged at 800rpm for 10min and then dissociated into a single cell suspension by using Trypsin/EDTA 0.05% and incubated 20min at 37°C. Primary cultures of spinal cord astrocytes were plated on poly-D-lysine (PDL) (Thermo Fisher Scientific, A3890401)-coated T25 flasks as described previously(Kempf, Knelles et al. 2021) Cells were grown in medium consisting of DMEM/F12 (1:1) with GlutaMax (Thermo Fisher Scientific, 35050061), 10% of FBS, 0.6% D-(+)-Glucose, 1% Penicillin/Streptomycin, 1x B27 (Thermo Fisher Scientific,12587010), 10ng/ml epidermal growth factor (EGF) (PreproTech, 100-47), and 10ng/ml basic fibroblast growth factor (bFGF). Cells were maintained in the incubator for 1 week at 37°C and 5% CO2. At 80%–90% confluency, cells were passaged using 0.25% trypsin/EDTA or frozen in freezing medium consisting of 60% Fetal Bovine Serum (FBS; Life Technologies, 26140-079), 30% medium and 10% DMSO (Sigma-Aldrich, C6295-50ML). Cells were fixed before the time of infection to evaluate the purity of the cultures.

For the *in vitro* infection of spinal cord astrocytes, the cells were thawed and plated at 20,000 cells/well in DMEM/F12 (1:1) GlutaMax, 10% of FBS, 0.6% D-(+)-Glucose, 1% Penicillin/Streptomycin, 1x B27, 10ng/ml EGF, and 10ng/ml bFGF onto 24 well plates (Nest Scientific, 70201) containing 25mm coverslips pre-coated with 1mg/ml of Poly-L-ornithine. 24 hours after plating, astrocytes were infected with 1.3µl of either Ascl1-mCherry (10^7^ - 10^8^ pfu) or miR24+miR9/9*+NeuroD1 (10^7^ - 10^8^pfu) suspended in DMEM with 10% FBS medium respectively. 16 hours after infection, the media was changed to fresh DMEM with 10% FBS and 5 μM Forskolin. From day 2 to day 5 post-infection, the media was replaced with DMEM/F12 (1:1) with N2 supplement, 10ng/ml bFGF, 200μM Ascorbic acid, 5μM Forskolin and 2ug/ml doxycycline. The last 10 days of reprogramming, the cells were maintained in the medium consisting of DMEM/F12 (1:1) with N2 supplement, 0.2M Ascorbic acid, 5μM Forskolin, 2μg/ml doxycycline, 20ng/ml BDNF, 20ng/ml GDNF, and 5μg/ml laminin added weekly into the medium.

### Immunocytochemistry

To determine the conversion efficiency of astrocytes into neuronal phenotypes *in vitro,* the cells were fixed with 4% paraformaldehyde (PFA) (Thermo Sci, C941G99) 2 weeks post-infection. For immunocytochemical characterization, cells were rehydrated for 15 minutes at room temperature in 1X phosphate-buffered saline (PBS). Fixed cells were quenched with the quenching solution, consisting of 30% methanol (Sigma-Aldrich, 646377-4L) and 2% hydrogen peroxide solution (H2O2) (EMD Millipore, HX06353) diluted in PBS for 1 hour at room temperature and then blocked with 10% Normal Donkey Serum (GeneTex, GTX73205), 0.3%Triton X-100 (Fischer Scientific, TX1568-1) in PBS for 1 hour at room temperature.

Cells were incubated with primary antibodies (Table 1) overnight (4°C), washed in PBS the following day (3x for 5 minutes), and incubated with secondary antibodies (Table 2) for 2 hours at room temperature. Immunolabelled cells were then washed, stained with 4,6-diamino-2-phenylindole (DAPI) (1:10,000; Invitrogen, D1306) in PBS for 10 minutes. Glass coverslips were mounted on microscope slides using Fluoro-Gel with Tris Buffer (Electron Microscopy Sciences, 17985–10). Cells were imaged and examined using a Zeiss inverted microscope (Zeiss Axio ImageM2 Apotome2). The efficiency of conversion was calculated as the percent of DAPI-positive cells that were also positive for the neuronal markers βIII-tubulin and MAP2.

**Table 1.** Primary antibodies.

| Antigen | Manufacturer | Host | Dilution | Catalog Number |
| --- | --- | --- | --- | --- |
| Anti- GFAP (polyclonal) | Abcam | Goat | 1:3000 | ab53554 |
| Anti-NeuN (monoclonal) | Millipore | Mouse | 1:1000 | MAB377 |
| Anti-MAP2 (monoclonal) | Abcam | Mouse | 1:500 | ab11267 |
| Anti- β-III tubulin (monoclonal) | Biolegend | Rabbit | 1:1000 | 802001 |
| Anti-TAUR1 (monoclonal) | donation from Dr Nicholas Kanaan, (Michigan State University) | Rabbit | 1:1000 | N/A |

**Table 2.**
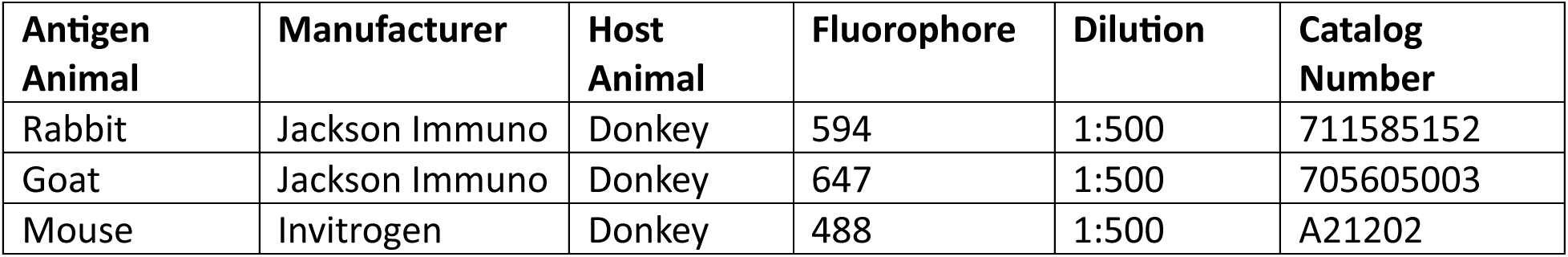
Secondary antibodies.

The diameter of nuclei (DAPI-positive cells), the diameter of cell bodies, and the length of the cellular processes in neurons were manually analyzed using Simple Neurite Tracer (SNT), an ImageJ plugin open-source tool. Analyses of neuronal cells were based on βIII-tubulin -positive cells and morphological parameters (e.g., presence of at least 2 processes longer than 3x the cell body). Analysis was performed by quantifying ∼50 neurons for each condition in 3 independent biological replicates.

### Neuronal activity analysis via Multi-Electrode Array (MEA) electrophysiology

To assess neurophysiological function, on day -2 of reprogramming, a subset of cultured brain and spinal cord astrocytes were replated onto Multiwell-MEA plates (Multi Channel Systems 60MEA200/30iR-Ti-gr). The wells were pre-coated overnight at room temperature with ∼0.07% polyethyleneimine (PEI) (Sigma-Aldrich, 181978) diluted in borate buffer [for 50ml: 155mg boric acid (Sigma-Aldrich, B0252), 237.5mg Borax (Sigma-Aldrich, B-9876), ddH2O to 50ml]. Plates were washed with ddH2O and then thoroughly dried for 1 h before plating cells. Cells were plated at a density of 50,000 cells per well and reprogrammed according to the protocol above. On day 6, cells were cultured in BrainPhys neuronal medium (Stem Cell Technologies, 05793) supplemented with ascorbic acid, forskolin, BDNF, GDNF, doxycycline, and weekly laminin. Spontaneous neuronal activity was recorded for 5 minutes using the Multiwell-Screen software (Multi Channel Systems MCS GmbH). Signals were sampled at 20kHz and filtered with a 350Hz fourth-order low-pass Butterworth filter and a 1Hz second-order high-pass Butterworth filter. Data analysis was performed using the Multiwell-Analyzer software (Multi Channel Systems MCS GmbH). Spikes were detected when the signal exceeded 6 standard deviations from the baseline noise. Network bursts were detected when there were six electrodes participating with three of them simultaneously participating. Spike and burst data were exported to Microsoft Excel for quantification. Neuronal activity was quantified.

Neuronal activity of brain-derived neurons was evaluated and confirmed pharmacologically by adding synaptic antagonists into the cell culture media. Activity was recorded before and 5–60 minutes after addition of NMDA receptor antagonist AP5 (50μM; Sigma-Aldrich, A8054), and before and 20–60 minutes after addition of GABA receptor antagonist bicuculline (10μM; Sigma-Aldrich).

Analysis of recordings focused on the following parameters: 1) the average number of active electrodes per well as a measure of consistent activity in cells throughout the well in each treatment condition; 2) the number of spikes at active electrodes per well as a measure of the amount of activity at each electrode under each condition, 3) the average number of electrodes with detectable bursting (at least one burst recorded), and the burst duration and the number of spikes per burst, as a measure of network activity. These parameters were used to measure activity under i) spontaneous and ii) pharmacologically modulated conditions using antagonists to glutamatergic and GABAergic receptors. Recordings were conducted 20-60 min after addition of NMDA receptor antagonist AP5 (50 μM) (Yates, Patil et al. 2021), and 20 min after addition of GABA receptor antagonist bicuculline (10 μM) (Dougherty, Bannatyne et al. 2005).

### Surgical Methods

Adult, female, Sprague Dawley rats (n=42) were housed at the Drexel University College of Medicine animal care facility. All experimental procedures were conducted with approval from the Institutional Animal Care and Use Committee and following the National Research Council Guidelines for the Care and Use of Laboratory Animals. Animals were anesthetized by injection of xylazine (10mg/kg, s.q.) and ketamine (120mg/kg, i.p.) for spinal cord injury (SCI), intraspinal virus injection and electrophysiology procedures.

#### Spinal Cord Injury (SCI)

For SCI, animals were prepared for surgery as previously described (Lane, Lee et al. 2012, Spruance, Zholudeva et al. 2018, Zholudeva, Iyer et al. 2018). Briefly, the back of the neck (from the base of the skull to the top of the shoulder blades) was shaved and sterilized with alternating alcohol wipes and iodine solution (ATC Medical, #233). Approximately 2.5 inch skin incision was made with a No. 15 scalpel blade from the base of the skull to the fifth cervical segment (C5), the surrounding musculature was cut and carefully pushed apart to expose the vertebral column. A laminectomy was made at the third cervical segment (C3) and rostral part of C4. Animals were suspended using surgical forceps secured at the C2 process and clamped at C4/5 vertebrae. Using the Infinite Horizon Pneumatic Impactor (Precision Systems, Lexington, Kentucky), with the impact force preset to 200 kilodynes and zero dwell time (actual force: 241.4 ± 12.9 kD; displacement: 1617.1 ± 33.5mm; velocity: 122.5 ±1.0mm/sec), animals received a lateralized contusion injury just caudal to C3 dorsal root on the left side of the spinal cord. All animals that went into respiratory arrest were intubated and placed on a mechanical ventilator (2.5ml, 70 breaths/minute; Small Animal Ventilator, Harvard Apparatus) for one hour. The underlying muscle was sutured (4-0 Vicryl Sutures, Fischer Scientific VCP310H) in layers and the skin was closed with wound clips (9mm; Fischer Scientific, # NC0142166).

#### Intraspinal Viral Injection

Injured animals were blindly separated into one of three groups (Control GFAP-mCherry, GFAP-Ascl1-mCherry and miR124+miR9/9*+NeuroD1-GFP) and decoded after all functional data analysis. Animals were anesthetized (xylazine-ketamine) as described above, the skin and underlying musculature were re-exposed one-week post-injury. Using a filling needle, 10µL of either control or active virus were preloaded into a 10µL Nanofil Syringe (World Precision Instruments, #NC0578818). The filling needle was replaced with a 35-gauge needle (World Precision Instruments, #NF35BV-2), and the syringe was secured to a micromanipulator (Model Kite-L, World Precision Instruments). Control (GFAP-mCherry, 10^7-^10^8^ pfu), Ascl1-mCherry (10^7-^ 10^8^ pfu), or miR-GFP (10^7-^10^8^ pfu), was injected into the contusion cavity (visualized by the bruise in the spinal tissue). The underlying muscle was sutured in layers and the skin was closed with wound clips as described above. Animals recovered for 8 weeks prior to terminal electrophysiology recordings.

#### Plethysmography

Whole body plethysmography (Data Science Instruments) was performed in awake, unrestrained animals to non-invasively measure ventilation 1 week prior to injury, 1 week after SCI, and weekly after viral injection, as previously described (Fuller, Doperalski et al. 2008, Lane, Lee et al. 2012). Briefly, animals were placed into the barometric plethysmography chambers with continuous air flow (breathing air) for 15-20 minutes to allow for acclimation and acquisition of baseline (eupneic) breathing. After acquiring baseline, the chambers were flushed with hypercapnic gas (7% CO_2_, balanced in N_2_) for 10 minutes to obtain ventilatory parameters while undergoing a respiratory challenge. Parameters which were quantitatively assessed from airflow traces included respiratory frequency (ƒ, breaths/min), tidal volume (V_T_ ml/100g) and minute ventilation (V_E_, ml/min/100g). Consistent with prior studies, hypercapnic challenge was employed to test the ability of the animals to increase ventilation upon demand. Data were averaged from the last 3 minutes during baseline acquisition and the last 3 minutes during hypercapnic challenge and reported as average value ± standard error. Statistical analyses were performed using SigmaStat 4.0 (Systat Software, Inc), with comparisons of each ventilatory parameter between each group and to pre-injury levels using repeated measures ANOVA and Student t-test.

#### Diaphragm Electromyography

Diaphragm activity was measured with terminal, bilateral electromyography (dEMG) recordings on every animal as described elsewhere (Lane, Lee et al. 2012, Zholudeva, Karliner et al. 2017, Spruance, Zholudeva et al. 2018), prior to being euthanized and perfusion-fixed for histological analyses. Briefly, animals were anesthetized with a mixture of xylazine and ketamine and a laparotomy was performed to expose the abdominal surface of the diaphragm. Bipolar hook electrodes (2 electrodes per side; PFA coated tungsten wire with exposed tips, A-M Systems, #796500) were then placed into the medial costal region of the left and right hemidiaphragm. Pulse oximetry was used to assess changes in blood oxygen saturation and estimate heart rate using a collar cuff (Part #: 015023) and MouseOx (Starr Life Sciences Corp.). Diaphragm activity was recorded during spontaneous breathing of room air and baseline was counted as a minimum of 10 minutes of stable activity with an oxygen saturation of >90%. After baseline was acquired, hypercapnic (7% CO_2_, balanced in N_2_; flow rate 2 L/min) or hypoxic gas (10% O_2_, balanced in N_2_; flow rate 2 L/min) was administered using a nose cone, as means of a 5-minute respiratory challenge. All raw dEMG signals were amplified (1,000x) and band pass filtered (0.3-10KHz) using differential A/C amplifier (Model 1700, A-M Systems) and digitized (Power 1401, Cambridge Electronic Design) until further analysis.

#### Immunohistochemistry of spinal cord tissues

At the end of terminal dEMG recordings, all animals were intracardially perfused with physiological saline (0.9% NaCl in water) and PFA (4% w/v in 0.1M PBS) pH 7.4. The brains and spinal cords of all animals were then removed and stored in 4% PFA overnight, at 4°C. Cervical spinal cord tissue spanning from cervical level 2 (C2) to just below C7 were blocked, cryoprotected (15%, then 30% sucrose, overnight at 4°C), embedded in M1 Embedding Matrix (ThermoFischer, #1310TS) and sectioned (on-slide 30µm, longitudinal) using a cryostat. To perform immunohistochemistry, sections were rehydrated for 15 minutes in PBS and blocked against endogenous peroxidase activity (30% methanol, 0.6% hydrogen peroxide in 0.1M PBS, incubated for 1 hour). Depending on the host species of secondary antibodies, sections were blocked against non-specific protein labeling (10% donkey or goat serum in 0.1M PBS with 0.02% Triton-X, incubated for 1 hour) prior to application of primary antibodies in blocking solution. Primary antibodies (summarized in Table 1) were left on the tissue overnight at in a humidified chamber at 4°C. The following day, the tissue was washed in PBS (3 times for 5 minutes each) and incubated in blocking solution with secondary antibodies for 2 hours (summarized in Table 2) at room temperature. Immunolabeled sections were then washed in PBS (3 x 5 minutes), allowed to dry, and coverslipped. Sections were examined using a Zeiss AxioImager microscope with Apotome 2, attached to a Dell PC. Images were taken with a digital camera (Zeiss Axiocam MRm).

## Statistical analysis

Cortical and spinal cord astrocytes-derived neurons: all data analyses were conducted blind and presented as mean ± SEM. At least three biological independent repeats were conducted for each experiment. Data were arranged in Microsoft Excel before statistical analyses were performed using Prism 10 (GraphPad). The statistical tests used were one-way analysis of variance (ANOVA). The Kruskal–Wallis test followed by Dunn’s multiple comparison test was used for data not normally distributed. Statistical significance was defined as *p<0.05, **p<0.005. The bar graphs reporting ‘ns’ were not significant according to non-parametric ANOVA tests.

<u>Rat functional electromyography:</u> raw diaphragm electromyography traces were integrated (DC bias removed at 0.1sec), rectified and smoothed (0.03sec) using Spike 2 software (version 8, Cambridge Electronic Design, UK). Baseline diaphragm output was determined during eupneic (room air) breathing by averaging the integrated signal over a 40 second interval immediately prior to a respiratory challenge. Diaphragm output during a respiratory challenge (i.e. during administration of hypoxic or hypercapnic gas) was determined by averaging integrated signal over a 40 second interval at the end of each of the 5-minute challenges. The average maximum voltage (the maximum height of the averaged breaths) and percent change of each challenge from baseline breathing was determined from the averaged, integrated signal for each respiratory state using OriginPro9 software (Northampton, Massachusetts). Results are reported as average maximum voltage +/-standard error, or as percent change relative to baseline (eupnea) where indicated. Statistical analyses were performed using SigmaStat 4.0 (Systat Software, Inc). Comparison between vehicle and transplant recipients was made using ANOVA and Student t-test. All analyses of electrophysiological data were blinded and decoded at the end of analysis.

